# Ablation of complex fractionated electrograms improves outcome in long standing persistent atrial fibrillation

**DOI:** 10.1101/123463

**Authors:** Claire A Martin, James P Curtain, Parag R Gajendragadkar, David A Begley, Simon P Fynn, Andrew A Grace, Patrick M Heck, Munmohan S Virdee, Sharad Agarwal

## Abstract

**Purpose:** There is controversy and sparse data on whether substrate based techniques in addition to pulmonary vein isolation (PVI) confer benefit in the catheter ablation of persistent atrial fibrillation (AF), especially if long standing. We performed an observational study to assess whether substrate based ablation improved freedom from atrial arrhythmia.

**Methods:** 286 patients undergoing first ablation procedures for persistent AF with PVI only, PVI plus linear ablation, or PVI plus complex fractionated electrogram (CFAE) and linear ablation were followed. Primary end point was freedom from atrial arrhythmia at one year.

**Results:** Mean duration of pre-procedure time in AF was 28+/-27 months. Freedom from atrial arrhythmia was higher with a PVI+CFAE+lines strategy then for PVI alone (HR 1.56, 95% CI: 1.04-2.34, p=0.032) but was not higher with PVI+lines. Benefit of substrate modification was conferred for pre-procedure times in AF of over 30 months. The occurrence of atrial tachycardia was higher when lines were added to the ablation strategy (HR 0.08, 95% CI: 0.01-0.59, p=0.014). Freedom from atrial arrhythmia at 1 year was higher with lower patient age, use of general anaesthetic (GA), normal or mildly dilated left atrium and decreasing time in AF.

**Conclusions:** In patients with long standing persistent AF of over 30 months duration, CFAE ablation resulted in improved freedom from atrial arrhythmia. Increased freedom from atrial arrhythmia occurs in patients who are younger and have smaller atria, and with GA procedures. Linear ablation did not improve outcome and resulted in a higher incidence of atrial tachycardia.

## Introduction

Atrial fibrillation (AF) is associated with increased mortality, congestive heart failure, and stroke. Catheter ablation (CA) has been established as a treatment option for patients with symptomatic, drug-refractory AF, with the best results obtained in patients with paroxysmal AF (PAF). In these patients, AF is usually related to the presence of triggers in the pulmonary veins; therefore, the generation of circumferential lesions around veins to electrically isolate them is the cornerstone treatment.

Results of CA in patients with persistent AF are less satisfactory [1]. Pulmonary vein isolation (PVI) is still central; however, recurrent and chronic AF induces progressive electrical and tissue structural remodeling, thus making AF a self-perpetuating disease [2]. To interrupt this process, addition of a substrate based ablation approach has been proposed; the two most common techniques used are linear lesions in the left atrium (LA) and focal ablation of complex fractionated atrial electrograms (CFAEs).

Linear lesions applied on the left atria roof and on the mitral isthmus are intended to modify the arrhythmogenic LA substrate and atrial macro-re-entrant circuits involved in maintenance of AF. Several studies have reported that these improve the efficacy of the procedure [3, 4]; however, if the lines are not electrically continuous, atrial tachycardias may develop. CFAEs are thought to indicate sites of conduction slowing or block, wavefront collision or anchor points for reentrant circuits. Several groups have reported improved outcomes using this technique in addition to PVI [5], but an important limitation is that CFAEs are largely identified by visual inspection and therefore dependent on operator judgement.

The STAR AF trial [6] in high-burden paroxysmal/persistent AF, showed that PVI plus CFAE ablation had the highest freedom from AF vs. PVI or CFAE ablation alone after one or two procedures. However, STAR-II AF [7], this time in solely persistent AF patients, contradicted these results, finding no difference in freedom for AF at 1 year between PVI alone, PVI + lines or PVI + CFAE ablation strategies, a finding subsequently replicated by other studies [8]. There is little data to support similar outcomes in long standing persistent AF

The HRS/EHRA/ECAS Expert Consensus Statement on Catheter and Surgical Ablation of Atrial Fibrillation from 2007 [9] states that ‘operators should consider more extensive ablation based on linear lesions or complex fractionated electrograms’ for ablation of persistent AF. However, in the light of recent evidence backing a PVI strategy alone [7, 10], we assessed the role of substrate modification in patients with long standing persistent AF. We also assessed the patient factors which could affect the outcome after ablation.

## Methods

### Study Design

This was an analysis of prospectively kept data of all cases of first catheter ablation for persistent AF ablation from the beginning of 2008 to the end of 2013 in a single large-volume centre. The study was approved by local ethics committee. Cases were excluded if they had had a previous ablation for either paroxysmal or persistent AF. All patients were over 18 years of age and all had symptomatic persistent AF (of over 7 days duration) refractory to at least one anti-arrhythmic agent. All patients were anticoagulated for at least 1 month pre-procedure and 3 months post-procedure, and given heparin to maintain an ACT of 300-350s during the procedure.

PVI was performed by either a point-by-point technique using Carto (Biosense Webster) or Ensite NavX (St Jude Medical) guidance or a single shot technique using a PVAC (Medtronic) or nMARQ (Biosense Webster) multi-polar catheter. The endpoint was PVI isolation confirmed by abolition of PV potentials. In patients undergoing substrate ablation, either linear ablation, CFAE ablation or both were undertaken using Carto or Ensite NavX guidance. Effective linear ablation was confirmed by bidirectional block across the roof of the LA and around the mitral valve isthmus tested by differential pacing techniques. CFAEs were defined as electrograms having a very short cycle length or fractionation composed of at least two deflections or perturbation of the baseline with continuous deflection of a prolonged activation complex [11]. CFAE detection was by visual inspection; automated software was not employed. CFAE ablation endpoint was the elimination of all CFAE sites identified in the LA and coronary sinus. Atrial tachycardias that occurred during the procedure were mapped and appropriate ablation performed. If the patients were in AF or an atrial tachycardia at the end of the procedure, they were cardioverted.

The strategy for ablation was operator based. The individual operators always followed a set strategy of either PVI only or PVI with substrate modification. The ablation strategy was not based on patient characteristics.

Data was collected of the patient demographics, of the ablation strategy and of the clinical outcome. Clinical assessments, 12 lead ECGs and 24 hr tapes were obtained at baseline and at 3, 6, 12 and 18 months after the ablation.

### Study outcomes

The primary study outcome was freedom from any documented episode of AF, at 1 year, lasting longer than 30s, occurring after the performance of first ablation procedures, with or without the use of anti-arrhythmic medications.

AF occurring within an initial 3 months blanking period after ablation was not counted, in accordance with the guidelines [9]. An episode of AF was considered part of the primary outcome analyses if it lasted longer than 30 seconds and was documented by any form of monitoring, regardless of symptoms. Patients who completed fewer than 3 months of follow-up and thus did not complete the blanking period were excluded from end-point analysis.

Main secondary outcomes included freedom from organised atrial arrhythmia (including atrial flutter or atrial tachycardia) after one ablation procedure and freedom from AF without use of antiarrhythmic medication.

### Statistical Analysis

We estimated a sample size of 80 per arm would be needed to provide 80% power (p < 0.05, 1 sided) to detect a 50% change in the hazard ratio (HR) between groups, assuming a median freedom from atrial arrhythmia of 12 months and a follow-up of 18 months [12].

Continuous variables are presented as mean ± standard deviation, and categorical data as counts or percentages. Analysis and comparisons of continuous data were performed using ANOVA, whilst the *χ*^2^ test was used to compare categorical data. Survival was estimated using Kaplan Meier (KM) analyses. Cox proportional-hazards-models were used to explore univariate and multivariate predictors of events. Initial exploratory co-variates of age, gender, LA diameter, left ventricular ejection fraction (LVEF), presence of risk factors including hypertension, diabetes, stroke, vascular disease and congestive heart failure (CHF), CHA_2_DS_2_-VASc score, use of general anaesthesia (GA) vs conscious sedation, type of mapping technique, length of time in AF, and use of anti-arrhythmic drug post procedure were used. Multivariate models included terms with *p*-value of <0.1 at univariate analysis. A two-sided probability level of <0.05 was considered significant. All calculations were performed using SPSS 20.0 (IBM Software, USA).

## Results

### Patients

A total of 333 patients were identified who had first ablations for persistent AF between 2008 and 2013. Of these 47 were excluded from the study because of: a lack of information regarding their procedure or follow-up (19), the procedure being abandoned before any ablation took place (5), the patients not in fact having the procedure they were labelled as having (6), the patients having had a prior PAF ablation (17).

Out of the 286 included patients, 79 had PVI alone, 85 had PVI plus lines, 15 had PVI plus CFAEs and 107 had PVI plus lines plus CFAEs. Lines included a roof line in all patients and an additional mitral isthmus line in 91 out of 192 (47%). Due to the relatively small number of patients undergoing PVI plus CFAEs, this group was excluded from further analysis as comparisons would not be statistically useful. Baseline characteristics were not significantly different between groups except LA diameter (χ^2^ = 0.034) (Table 1), where the proportion of patients with a severely dilated LA was higher in the those receiving PVI + CFAEs + lines than in the other groups.

**Table 1.** Characteristics of patients included in the study. Plus–minus values are means ±SD.

|  |  | **PVI alone** | **PVI + lines** | **PVI + CFAEs** | **PVI + lines + CFAEs** |
| --- | --- | --- | --- | --- | --- |
| Characteristic | Number | 79 | 85 | 15 | 107 |
|  | Age - yr | 59 +/- 9 | 59 +/- 10 | 59 +/- 7 | 59 +/- 9 |
|  | Male sex - no (%) | 60 (76) | 71 (84) | 11 (73) | 95 (89) |
|  | Procedure time - min | 176 +/- 38 | 186 +/- 37 | 183 +/- 27 | 191 +/- 39 |
|  | Fluoroscopy time - min | 39 +/- 19 | 41 +/- 25 | 39 +/- 26 | 36 +/- 24 |
|  | Ablation time - s | 2015 +/- 1333 | 2716 +/- 1224 | 2173 +/- 897 | 2771 +/- 1446 |
|  | GA - no (%) | 12 (15) | 9 (11) | 3 (20) | 17 (16) |
|  | Time in persistent AF - months | 30 +/- 27 | 27 +/- 26 | 22 +/- 13 | 28 +/- 29 |
| Medical history - no (%) | Hypertension | 31 (39) | 33 (39) | 3 (20) | 34 (32) |
|  | Diabetes | 1 (1) | 8 (9) | 0 (0) | 9 (8) |
|  | Vascular disease | 9 (11) | 14 (16) | 2 (13) | 9 (8) |
|  | Stroke/TIA | 3 (4) | 7 (8) | 0 (0) | 9 (8) |
|  | Heart failure | 9 (11) | 5 (6) | 0 (0) | 9 (8) |
| CHA<sub>2</sub>DS<sub>2</sub>-VASc score - n (%) | 0 | 35 (44) | 30 (35) | 10 (67) | 48 (45) |
|  | 1 | 29 (37) | 24 (28) | 5 (33) | 29 (27) |
|  | 2 | 11 (14) | 19 (22) | 0 (0) | 16 (15) |
|  | >2 | 4 (5) | 12 (15) | 0 (0) | 14 (13) |
| Baseline medications - n (%) | Beta-blocker | 34 (43) | 41 (48) | 4 (27) | 39 (36) |
|  | Ca channel blocker | 7 (9) | 5 (6) | 1 (7) | 8 (7) |
|  | Flecainide | 23 (29) | 19 (22) | 3 (20) | 23 (21) |
|  | Amiodarone | 16 (20) | 14 (16) | 0 (0) | 14 (13) |
|  | Dronedarone | 2 (3) | 0 (0) | 0 (0) | 2 (2) |
|  | Sotalolol | 7 (9) | 8 (9) | 2 (13) | 16 (15) |
| LA diameter - n (%) | normal | 45 (57) | 41 (48) | 8 (53) | 51 (48) |
|  | mildly dilated | 21 (27) | 26 (31) | 2 (13) | 30 (28) |
|  | moderately dilated | 11 (14) | 16 (19) | 5 (33) | 13 (12) |
|  | severely dilated | 2 (3) | 2 (2) | 0 (0) | 13 (12)* |
| EF - n (%) | normal | 62 (79) | 59 (69) | 13 (87) | 76 (71) |
|  | mildly impaired | 10 (13) | 11 (13) | 0 (0) | 16 (15) |
|  | moderately impaired | 4 (5) | 11 (13) | 0 (0) | 10 (9) |
|  | severely impaired | 3 (4) | 4 (5) | 2 (13) | 5 (5) |

\* denotes significant difference between groups (see main text)

A follow-up period of up to 18 months was examined. All patients had clinical assessment and 12 lead ECGs performed at each clinic visit. If there was no documentation of atrial arrhythmia but the patient described any symptoms of palpitations, SOB or fatigue which might be attributable to a recurrence of AF or to an organised atrial arrhythmia, they underwent Holter monitoring to achieve ECG documentation of a symptomatic episode (occurred in 22% of total patients).

### Primary outcome

At 12 months, freedom from atrial arrhythmia lasting longer than 30s after one ablation procedure, with or without the use of antiarrhythmic medications, was present in 29% of patients who underwent pulmonary vein isolation alone, 38% who underwent isolation plus lines, and 46% who underwent isolation plus lines plus CFAEs (Table 2).

**Table 2.** Outcome as a function of type of procedure

|  | **PVI alone**<br>**(n = 79)** | **PVI + lines (n**<br>**= 85)** | **PVI + CFAEs**<br>**(n = 15)** | **PVI + lines +**<br>**CFAEs (n = 107)** |
| --- | --- | --- | --- | --- |
| Freedom from any atrial<br>arrhythmia at 1 yr (%) | 27 (29) | 29 (38) | 8 (41) | 41 (46) |
| Freedom from atrial flutter<br>or tachycardia at 1 yr (%) | 67 (97) | 61 (84) | 14 (93) | 75 (83) |

There was no difference in freedom from atrial arrhythmia between PVI alone vs PVI plus lines (p = 0.879). The freedom from atrial arrhythmia was higher with a PVI plus CFAE plus lines strategy then for a PVI plus lines strategy (HR 1.51, 95% CI: 1.01 to 2.25, p = 0.043) or PVI alone (HR 1.56, 95% CI: 1.04 to 2.34, p = 0.032) (Figure 1).

**Figure 1.**
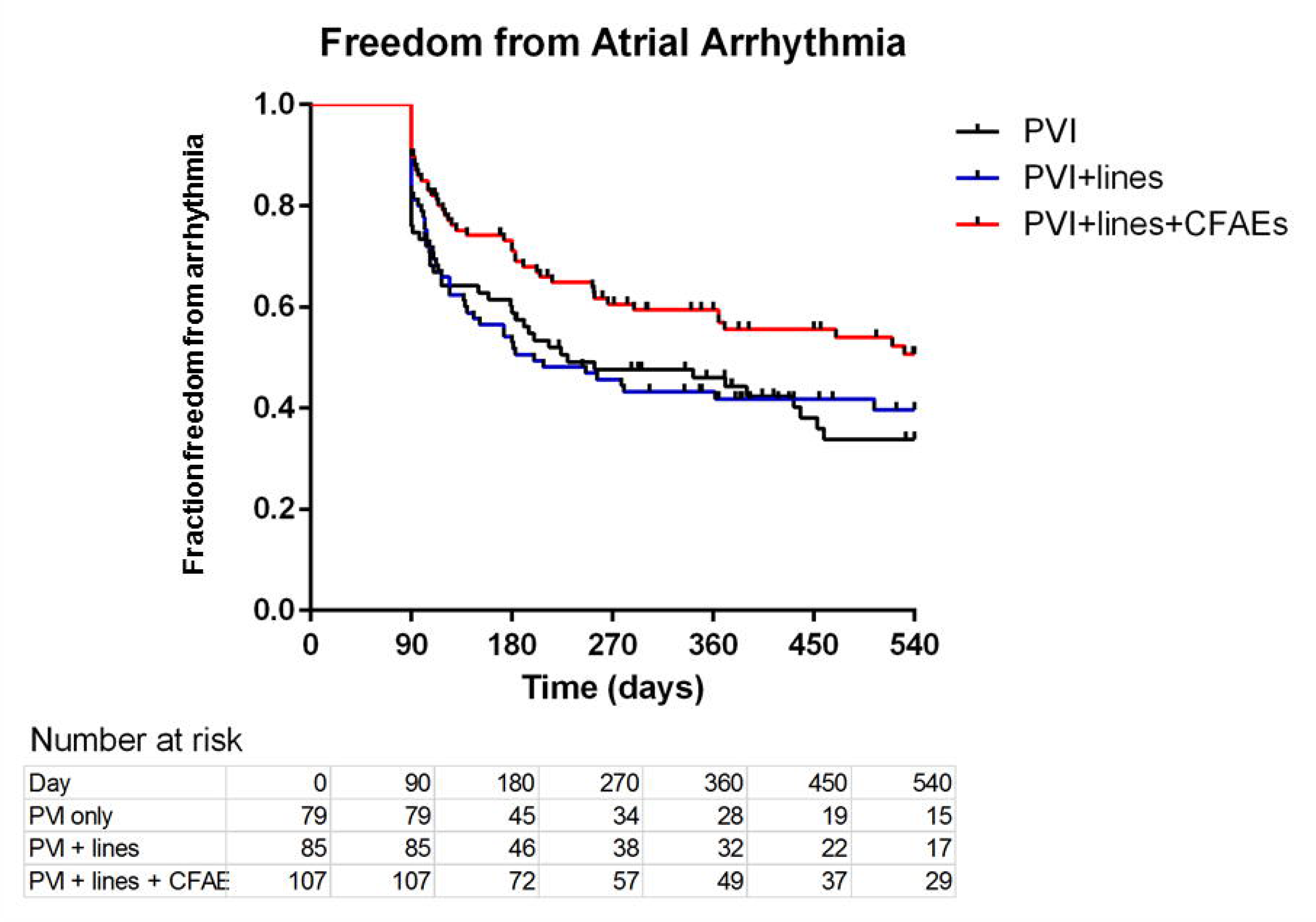
Kaplan Meier survival curves depicting freedom from atrial arrhythmia recurrence after one ablation procedure.

### Secondary outcomes

The freedom from atrial flutter or tachycardia was lower with a PVI plus lines strategy (HR 0.080, 95% CI: 0.01 to 0.59, p = 0.014) or a PVI plus lines plus CFAEs strategy (HR 0.078, 95% CI: 0.01 to 0.59, p = 0.013) than for PVI alone (Table2; Figure 2). There was no difference in outcome between PVI + Lines vs PVI + CFAE + Lines (HR 0.97, 95% CI: 0.49 to 2.02, p = 0.941).

**Figure 2.**
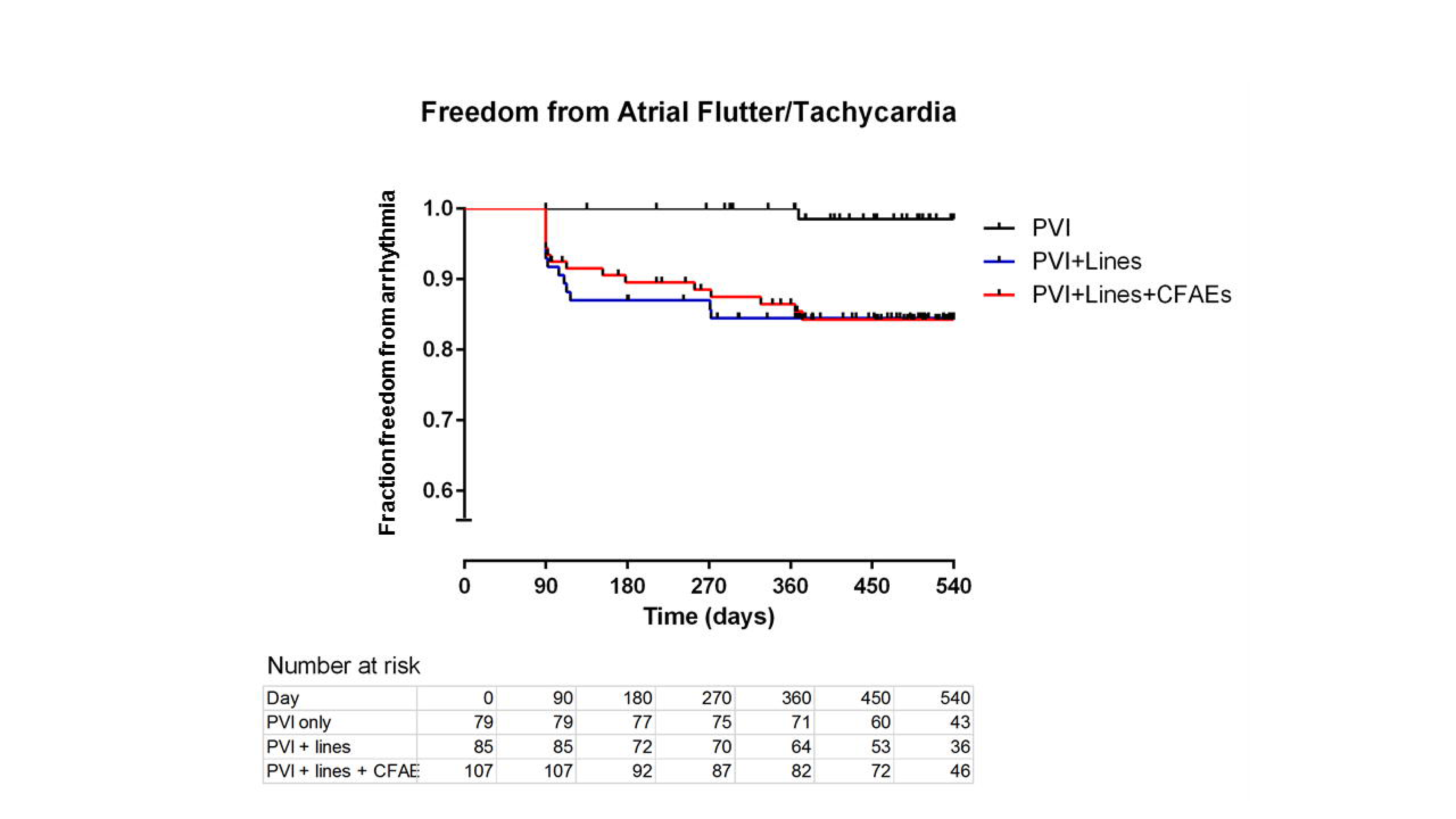
Kaplan Meier survival curves depicting freedom from occurrence of organised atrial arrhythmia (atrial flutter or atrial tachycardia) after one ablation procedure for persistent AF.

### Adverse events

1 groin haematoma occurred in patients undergoing PVI only; 1 tamponade and 1 stroke occurred in patients undergoing a PVI plus lines strategy; and 2 tamponades, 1 stroke, 1 groin haematoma and 1 reactive pericardial effusion occurred in patients undergoing a PVI + CFAEs + lines strategy. There were a further 5 cases that were hindered by problems with sedation, with patients becoming agitated or with uncontrolled pain; these were split between PVI alone (2), PVI plus lines (1) and PVI plus CFAEs (2).

### Subgroup analysis

Freedom from any atrial arrhythmia and freedom from atrial flutter or tachycardia was analysed stratified by age, gender, whether GA was used, presence of risk factors including stroke, diabetes, CHF, hypertension and PVD, CHA_2_DS_2_-VASc score, LA diameter, EF, time in AF, and whether an anti-arrhythmic (amiodarone, dronedarone, flecainide or sotolol) was used post procedure.

Univariate Cox regression analysis found the following factors to increase likelihood of freedom from atrial arrhythmia at the p < 0.1 level (Table 3): decreasing age, use of GA over conscious sedation, normal or only mildly dilated LA, decreasing time in AF pre-procedure, use of a ‘point-by-point’ rather than ‘one-shot’ technique and use of an antiarrhythmic drug post procedure. In multivariate analysis, there was higher freedom from atrial arrhythmia with a PVI plus CFAE plus lines strategy then for a PVI alone, as well as for decreasing age, use of GA over conscious sedation, normal or only mildly dilated LA and decreasing time in AF pre-procedure.

**Table 3.** Factors influencing freedom from atrial arrhythmia at 1 year. Bold text denotes significant difference between groups (see main text)

|  | Univariate |  |  | Multivariate |  |  |
| --- | --- | --- | --- | --- | --- | --- |
| Predictors | HR | 95% CI | p-value | HR | 95% CI | p-value |
| Age /year | 0.983 | 0.996 - 1.000 | **0.051** | 0.960 | 0.96 - 0.99 | **0.012** |
| Gender - male | 0.797 | 0.542 - 1.173 | 0.250 |  |  |  |
| Normal/mildly dilated LA | 1.460 | 1.017 - 2.083 | **0.033** | 1.510 | 1.020 – 2.273 | **0.041** |
| Normal LV function | 1.170 | 0.851 – 1.608 | 0.334 |  |  |  |
| Hypertension | 1.054 | 0.683 - 1.32 | 0.756 |  |  |  |
| Diabetes | 0.884 | 0.577 - 2.218 | 0.721 |  |  |  |
| Stroke | 0.706 | 0.663 - 3.024 | 0.366 |  |  |  |
| Vascular disease | 0.604 | 0.540 - 3.704 | 0.386 |  |  |  |
| CHF | 1.302 | 0.451 - 1.31 | 0.333 |  |  |  |
| CHA2DS2-VASc score $\geq 2$ | 0.807 | 0.559 – 1.166 | 0.254 | | | |
| Use of GA | 1.721 | 0.378 – 0.892 | **0.013** | 1.600 | 1.01 - 2.55 | **0.048** |
| Ablation technique – ‘point by point’ vs ‘one-shot’ | 1.486 | 1.053 - 2.096 | **0.024** | 1.404 | 0.973 – 2.028 | 0.070 |
| Length of time in AF/months | 0.995 | 0.999 - 1.011 | **0.086** | 0.995 | 0.990 - 1.000 | **0.041** |
| Anti-arrhythmic post-procedure | 1.039 | 0.994 – 1.086 | **0.091** | 1.287 | 0.814 – 2.033 | 0.280 |
| PVI+lines vs PVI only | 1.031 | 0.693 – 1.443 | 0.879 |  |  |  |
| PVI+CFAE+lines vs PVI only | 1.558 | 1.038 – 2.336 | **0.032** | 1.610 | 1.04 - 2.49 | **0.022** |
| PVI+CFAE+lines vs PVI+lines | 1.508 | 1.012 – 2.247 | **0.043** | 1.374 | 0.881 – 2.141 | 0.161 |

In univariate analysis the following factors increased the likelihood of freedom from atrial flutter or tachycardia at the p < 0.1 level (Table 4): use of GA over conscious sedation, use of a ‘point-by-point’ rather than ‘one-shot’ technique, absence of hypertension and CHA_2_DS_2_-VASc score of 0 or 1. In multivariate analysis, the freedom from atrial flutter or tachycardia was higher with a PVI only strategy than for PVI plus lines or PVI plus lines plus CFAEs, as well as for use of GA over conscious sedation.

**Table 4.** Factors influencing freedom from atrial flutter or tachycardia at 1 year. Bold text denotes significant difference between groups (see main text)

|  | Univariate |  |  | Multivariate |  |  |
| --- | --- | --- | --- | --- | --- | --- |
| Predictors | HR | 95% CI | p-value | HR | 95% CI | p-value |
| Age/year | 0.980 | 0.942 – 1.021 | 0.348 |  |  |  |
| Gender - male | 0.622 | 0.218 - 1.779 | 0.372 |  |  |  |
| Normal/mildly dilated LA | 1.018 | 0.439 – 2.364 | 0.967 |  |  |  |
| Normal LV function | 1.248 | 0.615 – 2.351 | 0.540 |  |  |  |
| Hypertension | 0.439 | 0.216 - 0.891 | **0.023** | 0.634 | 0.222 – 1.812 | 0.395 |
| Diabetes | 0.452 | 0.158 – 1.292 | 0.139 |  |  |  |
| Stroke | 0.997 | 0.238 - 4.177 | 0.996 |  |  |  |
| Vascular disease | 0.844 | 0.633 – 2.143 | 0.886 |  |  |  |
| CHF | 0.819 | 0.249 - 2.693 | 0.741 |  |  |  |
| CHA2DS2-VASc score $\geq 2$ | 0.314 | 0.155 – 0.635 | **0.001** | 0.571 | 0.150 - 2.169 | 0.410 |
| Use of GA | 2.268 | 1.101 – 4.673 | **0.026** | 3.125 | 1.076 – 9.091 | **0.036** |
| Ablation technique – ‘point by point’ vs ‘one-shot’ | 1.854 | 0.917 – 3.751 | **0.086** | 1.759 | 0.863 – 3.586 | 0.120 |
| Length of time in AF/months | 1.000 | 0.987 - 1.014 | 0.915 |  |  |  |
| Anti-arrhythmic post-procedure | 1.139 | 0.558 - 2.325 | 0.720 |  |  |  |
| PVI+lines vs PVI only | 0.077 | 0.010 – 0.592 | **0.014** | 0.160 | 0.340 - 0.710 | **0.016** |
| PVI+lines+CFAEs vs PVI only | 0.078 | 0.010 – 0.587 | **0.013** | 0.074 | 0.010 – 0.562 | **0.012** |
| PVI+CFAE+lines vs PVI+lines | 0.973 | 0.486 – 2.022 | 0.941 |  |  |  |

The time in AF pre-procedure was far greater in our patients (mean 28 +*/-* 27 months) than in most studies examining catheter ablation of persistent AF [7]. In order to examine this more closely, KM curves were generated stratified into patients with long-standing persistent AF, defined as persistent AF for over 30 months (n = 112), vs the rest. This demonstrated that there was no significant benefit in additional ablation techniques over PVI isolation in those who had AF for under 30 months (p = 0.295). However, the benefit in PVI + CFAE + lines ablation persisted for those for had had AF for over 30 months preprocedure HR = 1.82, 95% CI 1.00 to 3.33, p = 0.041) (Figure 3).

**Figure 3.**
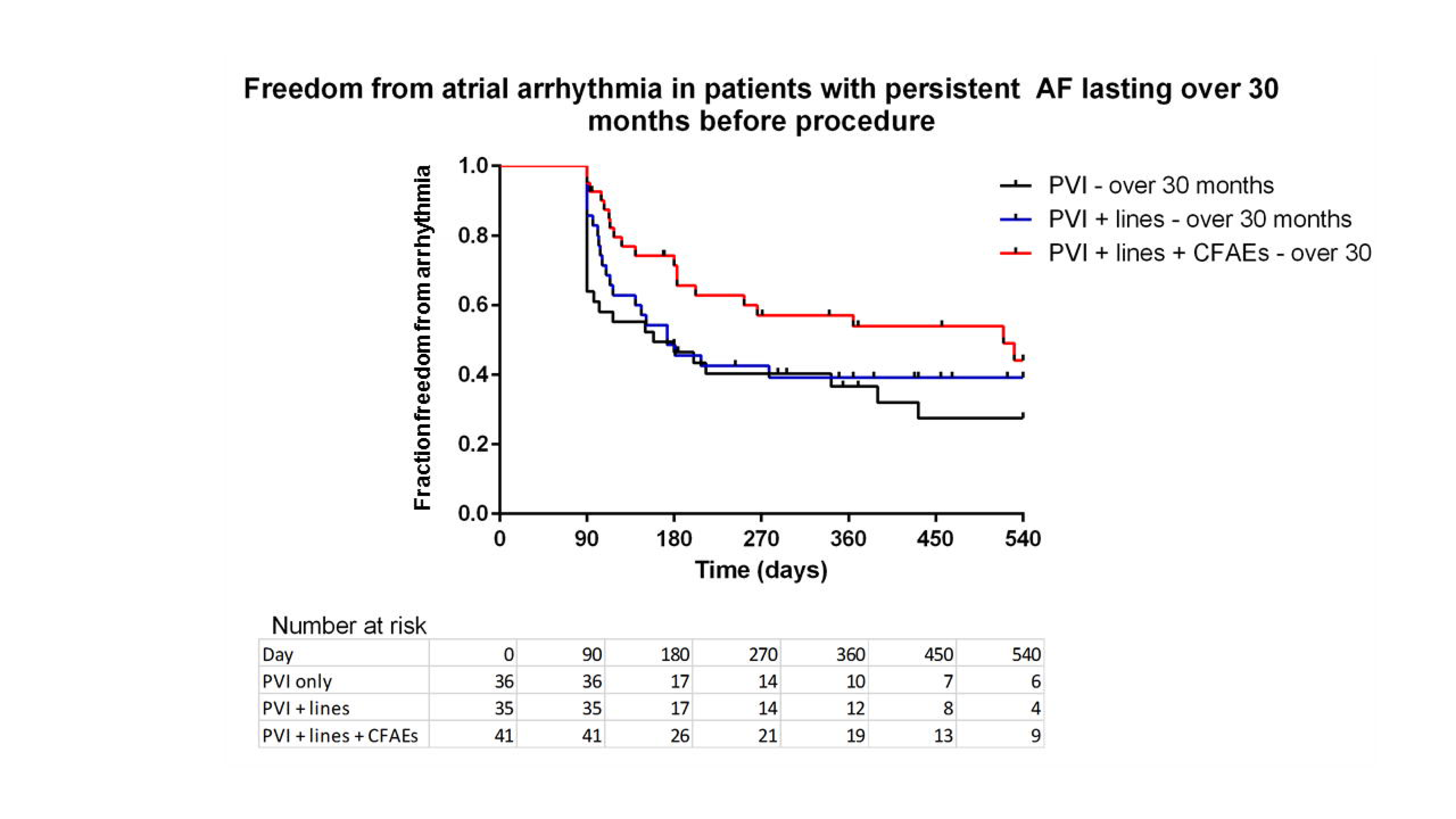
Kaplan Meier survival curves depicting freedom from atrial arrhythmia after one ablation procedure in patients with persistent AF lasting over 30 months before their procedure

## Discussion

This is an observational study of persistent AF ablation in a single large-volume centre. The majority of patients had long standing persistent AF by the time they came for ablation. Freedom from atrial arrhythmia was higher with a PVI plus CFAE plus lines strategy than for a PVI plus lines strategy or PVI alone, suggesting a possible role for CFAE ablation in addition to PVI but not linear ablation. On sub-analysis, this effect was only significant for patients that had been in AF for over 30 months.

It is interesting that these results contradict those of Star-II AF and some recent studies [7, 8, 13]. The reason is likely to lie in the differing demographic of the patient population, especially the duration of AF prior to the ablation. In the Star-II AF trial, only 70-80% were in persistent AF for over 6 months, and those in PAF had a relatively low AF burden of 80-85 hours/month. 95% of the patients in our study had been in persistent AF for over 6 months, and the mean duration of persistent AF was 28 +/− 27 months. AF is perpetuated by electroanatomic remodelling which develops as AF progresses over time. This may involve atrial dilatation, interstitial fibrosis, uncoupling and loss of myofibrils, deposition of extracellular matrix, loss of gap junctions, and resultant anisotropy and conduction slowing and block and refractory period heterogeneity. Therefore in patients with long standing persistent AF, abnormal substrate may have a stronger role to play in AF maintenance than simple triggers targeted primarily by PVI, and a substrate based approach may be of benefit. Our study also demonstrated that the addition of linear ablation results in a higher propensity to post-procedure organised atrial arrhythmias. The incidence of post-procedure atrial flutter or tachycardia was not significantly worse with CFAE ablation plus lines plus PVI over PVI plus lines.

We have demonstrated worse outcomes in terms of freedom from AF with increasing age, duration of persistent AF and LA size. There is a lack of randomized control trials focusing on the elderly group to measure the outcomes of catheter ablation. Single centre studies mostly examining PAF ablation have not demonstrated a significant difference in outcome with age [14, 15], although a higher percentage of patients required antiarrhythmic drugs to maintain sinus rhythm [15]. It is recognised that diastolic LV dysfunction, a condition that is well-linked with aging, leads to increased atrial filling pressures and wall stretch, ultimately associated with atrial fibrosis and remodelling, which make AF more likely to persist [16]. Our study suggests that elderly patients with long standing persistent AF are less likely to gain benefit.

It has been recognised for some time that increased LA size is a predictor for AF recurrence [17]. Specifically, our study would suggest that targeting ablation for patients with normal or mildly dilated LAs would be more effective. We did not find any significant deterioration in outcome with increasing co-morbidities or CHA_2_DS_2_-VASc score.

There was initially benefit from the addition of an anti-arrhythmic drug post-procedure, but this did not confer long term benefit. Previous studies in paroxysmal AF have suggested the addition of anti-arrhythmics are beneficial in the first weeks following ablation but have failed to demonstrate long term benefit [18, 19].

We found that outcome of ablation is improved if the procedure had been conducted under GA rather than conscious sedation. There is emerging evidence that GA may improve outcome, whilst reducing fluoroscopy and procedure times [20]. This may be due to patient immobility, improved accuracy of mapping, and catheter stability, allowing better contact between the catheter and the endocardium and optimizing lesion quality. There has been concern that atrio-oesophageal fistula may be more common when the procedure is performed under GA [21]; no instances occurred in our study in either group.

The main limitations of this study lie in it being a retrospective analysis. Ablation technique was operator dependent, where some operators preferentially did only PVI and others did additional substrate modification as their standard index procedure. Ablation strategy however, was not based therefore on individual patient factors. There were insufficient number of patients in the PVI + CFAE arm to provide adequate power for analysis of the group in a meaningful way to the other groups. In some areas it would have been useful to have more robust data collection, for example on adverse events, particularly for vascular access complications. Continuous monitoring was not employed to monitor for possible recurrence of paroxysmal AF. Furthermore, it would have been useful to gather Quality of Life information from patients undergoing AF ablation to better analyse the impact of ablation of the patient. These results are nonetheless hypothesis generating and may warrant a randomized trial for patients with more long-standing arrhythmia.

## Conclusion

Based on our findings, we would suggest consideration of the addition of CFAE ablation to PVI in patients with long standing persistent AF, especially in those with persistent AF for over 30 months. We have not demonstrated any benefit in linear ablation over PVI alone, and linear ablation resulted in a higher incidence of organised atrial arrhythmia. In selecting patients who might derive most benefit from persistent AF ablation, we would suggest selecting those who are younger and with smaller LA. We would recommend the use of GA over conscious sedation. In our region, there is a significant time from onset of AF to ablation. This is due to a variety of factors, which could be potentially reduced by educating patients so that a diagnosis is reached quicker, direct referral to electrophysiologists, and reducing clinic and procedure waiting times.

## Compliance with ethical standards

### Conflict of Interest

The authors declare that they have no conflict of interest.

All procedures performed in studies involving human participants were in accordance with the ethical standards of the institutional and with the 1964 Helsinki declaration and its later amendments. For this type of retrospective study formal consent is not required.

